# AntiCP3: Prediction of Anticancer Proteins Using Evolutionary Information from Protein Language Models

**DOI:** 10.1101/2025.04.29.651196

**Authors:** Amisha Gupta, Milind Chauhan, Ritu Tomer, G.P.S. Raghava

## Abstract

A number of computational methods have been developed in the past for predicting anticancer peptides, including AntiCP and AntiCP2 from our group. While these tools have been widely used by the scientific community, they are not suitable for predicting anticancer proteins. In this study, we present AntiCP3, the first dedicated method for the prediction of anticancer proteins. All models were trained using five-fold cross-validation and evaluated on an independent dataset not used during training. Our initial analysis revealed distinct compositional differences between anticancer peptides and proteins, justifying the need for a separate prediction framework. We first implemented similarity-based approaches, which yielded moderate performance. Subsequently, we developed machine learning and deep learning models using conventional protein features, achieving a maximum AUC of 0.72. The performance improved to an AUC of 0.79 with the incorporation of evolutionary information through PSSM profiles. Further enhancement was observed when embeddings from a fine-tuned protein language model ESM-t33 were used, leading to a best AUC of 0.90. Finally, a hybrid approach combining BLAST with our machine learning model achieved an AUC of 0.91. To facilitate the scientific community, we have implemented AntiCP3 as both a web server and standalone software for the prediction of anticancer proteins (https://webs.iiitd.edu.in/raghava/anticp3/). We have also deployed our model at hugging face https://huggingface.co/raghavagps-group/anticp3.

**Highlights:**

⍰ Existing methods have been developed for predicting anticancer peptides.
⍰ AntiCP3 is specifically optimised for predicting anticancer proteins.
⍰ PSI-BLAST used to obtain evolutionary information in form of PSSM profile.
⍰ Hybrid methods developed using alignment free and alignment based approach.
⍰ A web server and standalone tool have been developed to assist the community.

## Introduction

Cancer is one of the most devastating group of disease which leads to estimated 611,720 deaths in US, in 2024 (“Erratum to ‘Cancer Statistics, 2024’.,” 2024). It arises due to the uncontrolled growth of cells, often developing over an extended period (Ganesh & Massague, 2021). Various therapeutic approaches have been designed to combat this life threatening disease such as traditional treatment (surgery, chemotherapy and /or radiotherapy), advanced strategies such as exosome based therapy, stem cell based therapy, gene therapy, personalized medicines (Kaur et al., 2023). Despite advancement in diagnosis and treatment, it is still a major health concern across the globe. Now-a-days, protein based therapeutics have emerged for the treatment purposes of various diseases like cancer, autoimmune diseases, neurodegenerative disorder and other (Baig et al., 2018; Jain, Gupta, Patiyal, et al., 2024; Usmani et al., 2017). Presently, numerous protein/peptides are being used for the therapeutic purposes such as anti-diabetic, anticancer, and antithrombotic protein/peptides (Baig et al., 2018). Till now, 894 proteins including 354 monoclonal antibodies (mABs) and 85 peptides or polypeptides were approved by FDA (Jain, Gupta, & Raghava, 2024; Jain, Gupta, Patiyal, et al., 2024; Usmani et al., 2017). FDA have already approved number of peptide-based anticancer drugs (e.g., Pembrolizumab, Dactinomycin, Plitidepsin) and many more are under in clinical trials (Chinnadurai et al., 2023; Mukherjee et al., 2023). CancerPPD-2 also contains around 47 entries of clinical trials for the anticancer peptides/proteins (Milind et al., 2025).

Both large proteins or short peptides have their own significance in therapeutic approaches. Small peptides are easy to synthesis, having high target specificity and selectivity with low toxicity but low stability and short half-life (Marqus et al., 2017). On the other hand, proteins have intricate structures that allow them to specifically bind to unique cancer cell markers such as mABs. This specificity minimizes off-target effects and reduces damage to healthy cells (Mukherjee et al., 2023; Wang et al., 2019). Anticancer proteins have number of advantage over traditional drugs in manging treatment of cancer patients. They can be utilised as immunotherapy, targeted therapy, DNA repair inhibition, angiogenesis inhibition, apoptosis induction and tumor suppression (Ashkenazi & Dixit, 1998; Lord & Ashworth, 2017; Mellman et al., 2011; Scott et al., 2012; Vogelstein et al., 2000; Vousden & Prives, 2009; Zou, 2005). In the past number of methods have been developed to predict anticancer peptides (Agrawal et al., 2021; Geng et al., 2025; Kilimci & Yalcin, 2024; Tyagi et al., 2013; Zhu et al., 2022).

The major focus of these methods is to predict anticancer activity of peptides or small proteins, mainly trained on peptides in CancerPPD (Tyagi et al., 2015). In this study, we have made a systematic attempt to develop anticancer activity of proteins. We have extracted anticancer proteins as positive dataset from CancerPPD-2.0 (Milind et al., 2025), and non-anticancer proteins from Uniprot for negative dataset using different keywords. Here, we have extract different set of features such as sequence based features (composition-based features), structure based feature (RSA, secstate), and evolutionary profile based features. Compositional based features helps in understanding specific amino acid contribution individually and in group (Meher et al., 2017), while evolutionary profile based feature helps in identifying the conserved functional regions of the given protein (Gil & Fiser, 2018). The secondary structure based features such as Secondary structure state (Secstate) and Relative solvent accessibility (RSA) provides the detailed information about percentage of residue in each secondary state in a protein and also about the relative solvent accessibility of each residue in 3D structure (Tien et al., 2013). We have also extract sequence embeddings using fine-tuned esm2 protein language models. These features were further used for the classification anticancer non-anticancer proteins using machine learning based classifiers. We have also applied alignment based approaches like BLAST and Merci motifs. We achieved best performance when we combined machine learning models alignment based methods called hybrid methods. Finally, we have incorporated best performing fine-tuned esm2 model as well as hybrid model in the webserver. We have also developed GitHub repository, pypi package, hugging face and python based standalone package.

## Material and Methodology

In this study we obtained anticancer proteins from CancerPPD2 and non-anticancer proteins from UniProt. All models have been trained, tested and evaluated on non-redundant set of proteins where no two proteins have more than 40% similarity with other proteins. The process of extracting data and creating non-redundant dataset is shown in **figure 1**.

**Figure 1:**
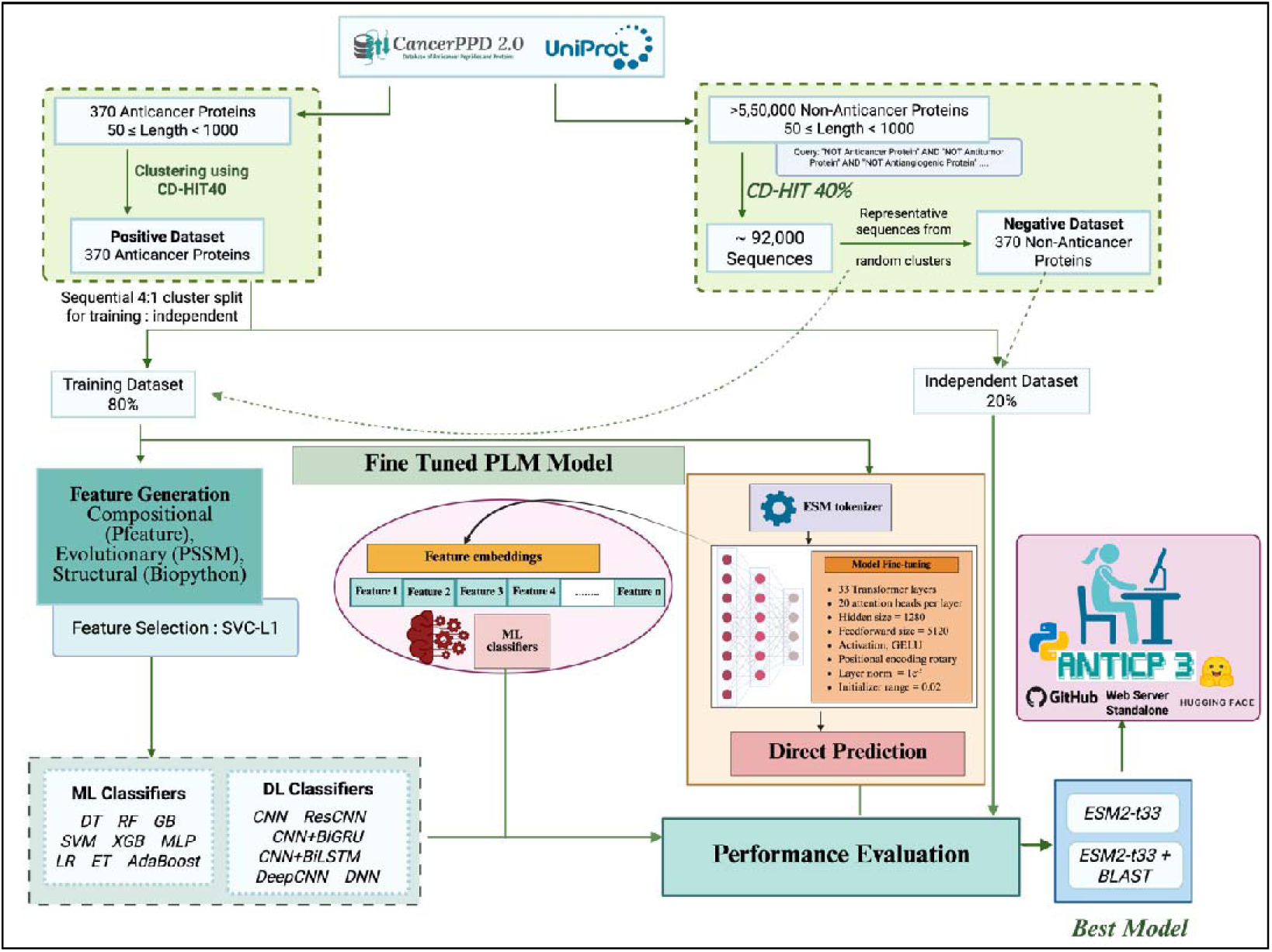
The figure depicts the overall architecture of the study.

1. Dataset Collection & Compilation We have collected the anticancer protein dataset from CancerPPD2 database, which contains anti-cancer peptides and proteins sequences. We have retrieve 370 natural protein sequences using length filter ranges between (50 to 1000). This dataset is named as Positive dataset for this study. For negative dataset, we have mined UniProt database using keywords “NOT - Anticancer protein”, “NOT - Antitumor protein”, and “NOT - Antiangiogenic protein” AND “length: 50 TO *”. We have retrieved total 5,58,717 sequences as non-anticancer proteins.
2. Data Preprocessing Data preprocessing is crucial step while performing classification task. Here, we have applied several preprocessing steps to clean any redundant or highly similar sequence in the positive or negative set. At first, we have applied CD-HIT40 on positive as well as on negative dataset to remove highly similar sequences as suggested in other studies (Khanduja & Mohanty, 2025; Li & Godzik, 2006; Sangaraju et al., 2024). In positive set, we have obtained 205 clusters after applying CD-HIT40 which were then used for further analysis. Here, we use all clusters for positive data as we have less amount of known anticancer proteins. The positive data splitting in training and validation can be seen in figure 2. In the given figure, we have explained fold splitting in two sets i.e. outer validation and inner validation. The outer validation means the data is splitting into main training fold and unseen validation fold. The inner validation means the training fold of outer validation is further splitting into training and internal validation process to overcome bias, variance and overfitting in performance evaluation. After training the model using internal validation, we reevaluate our model performance on outer validation which was never seen by the model before. Then we apply CD-HIT40 on negative set to remove highly similar sequences. We have selected only the representative sequence of each cluster after which 92,388 protein sequences were left. We have also confirmed that our both positive and negative dataset are unique and not having any redundant sequence. We have also checked for non-natural amino-acids present in protein sequences. If any sequence contains non-natural amino-acids then we removed those sequences from the final dataset. Then we randomly select 370 sequences with length ranges between 50-1000 for negative dataset as our positive dataset contains same length of sequences. Finally, we have 370 random proteins same as of anticancer proteins for classification purpose.
3. Feature Generation Extracting sequence-based features that capture structural, physicochemical, and compositional qualities is crucial to improving our understanding of anti-cancer proteins. In this study, we have extracted three different type of features using protein sequences. 1. Composition-based features 2. Secondary Structure-based features 3. Evolutionary Profile-based features 4. Sequence Embedding.

### 3.1. Composition-based features

We have computed composition-based features of protein sequences using the Pfeature tool. From the number of features offered by this tool, we have computed amino acid composition, di-peptide composition and physiochemical property based features for this study. To calculate the amino acid composition and di-peptide composition based features following formulas were used:

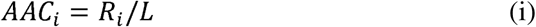

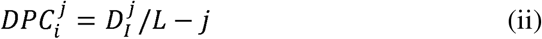

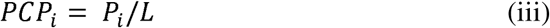

Where, AAC_i_ is amino acid composition of residue type i; R_i_ and L number of residues of type i and length of sequence, respectively. DPC^j^ is the fraction or composition of dipeptide of type i for jth order. D^j^ and L are the number of dipeptides of type i and length of a protein sequence, respectively., PCPi is Physico-chemical properties composition of type i; Pi and L are sum of property of type i, and length of sequence, respectively.

### 3.2. Secondary Structure based features

To calculate secondary structure based features from sequence, 3D structures of these proteins were required. Unfortunately, the experimentally validated 3D structure of anticancer proteins used in this study are not available. So, here we obtained the 3D structures of these proteins from UniProt which were predicted using Alpha fold server. These predicted structures were then utilized for feature extraction. Here, we used biopython’s DSSP module for structure based features extraction (Miller et al., 1987; Rost & Sander, 1994; Tien et al., 2013). This module is designed to extract Secondary structure state and Relative solvent accessibility (RSA) against each residue in a protein sequence.

The secondary structure state provided by DSSP module calculates H: Alpha helix (4-12); B: Isolated beta-bridge residue; E: Strand; G: 3-10 helix; I: Pi helix; T: Turn; S: Bend; -: None; P: PolyProline Helix II (Chakrabarti, 2025).

Further, we calculated the percentage of residues in each state in individual protein sequence and used these percentages as secondary state features in N*9 vector, where N refers to protein sequence. We have provided the example secondary state feature matrix in Figure 2, for better understanding of dataset.

**Figure 2:**
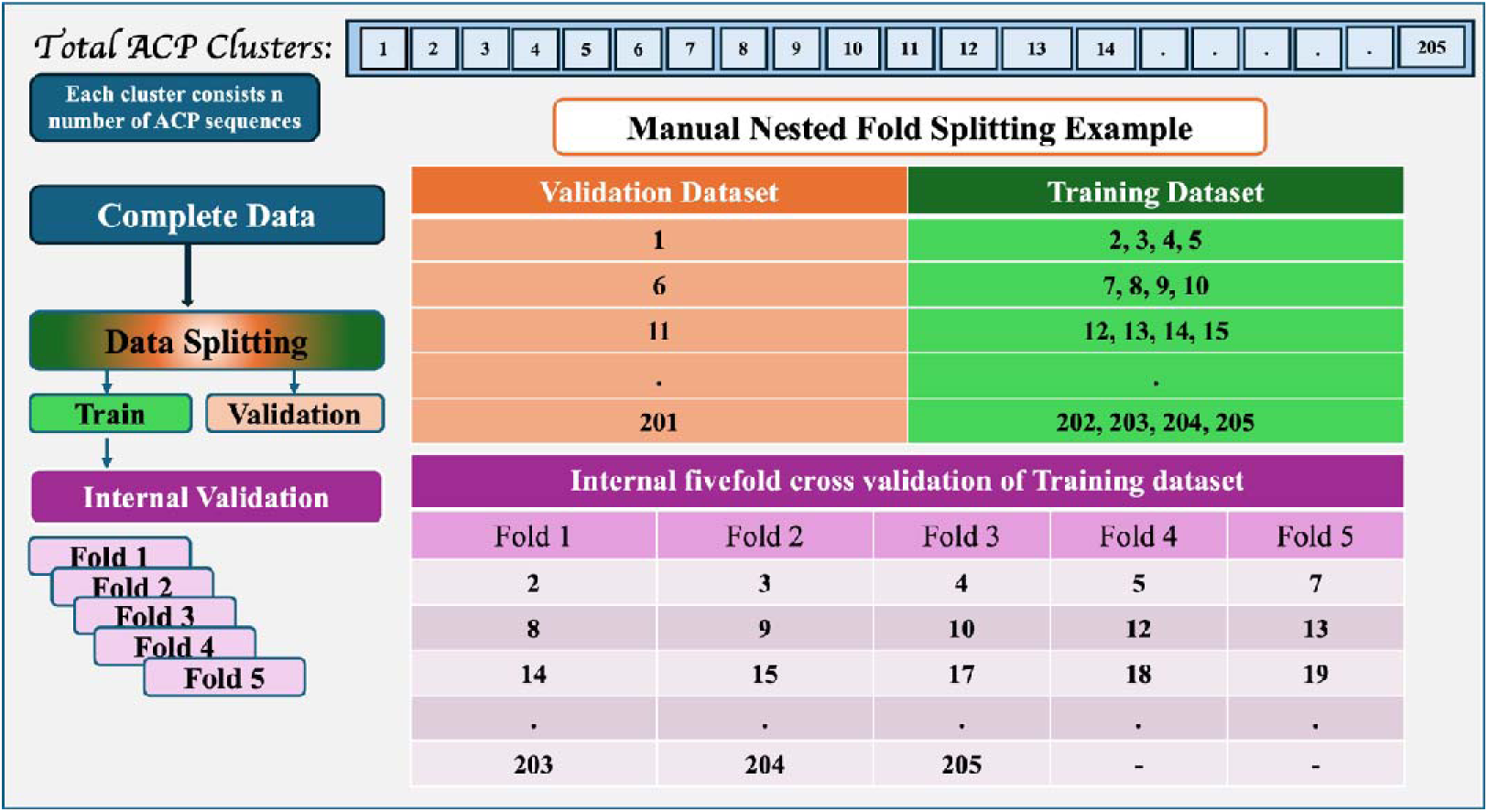
The figure shows the manual fold splitting of anticancer proteins for more robust model development.

**Figure 2:**
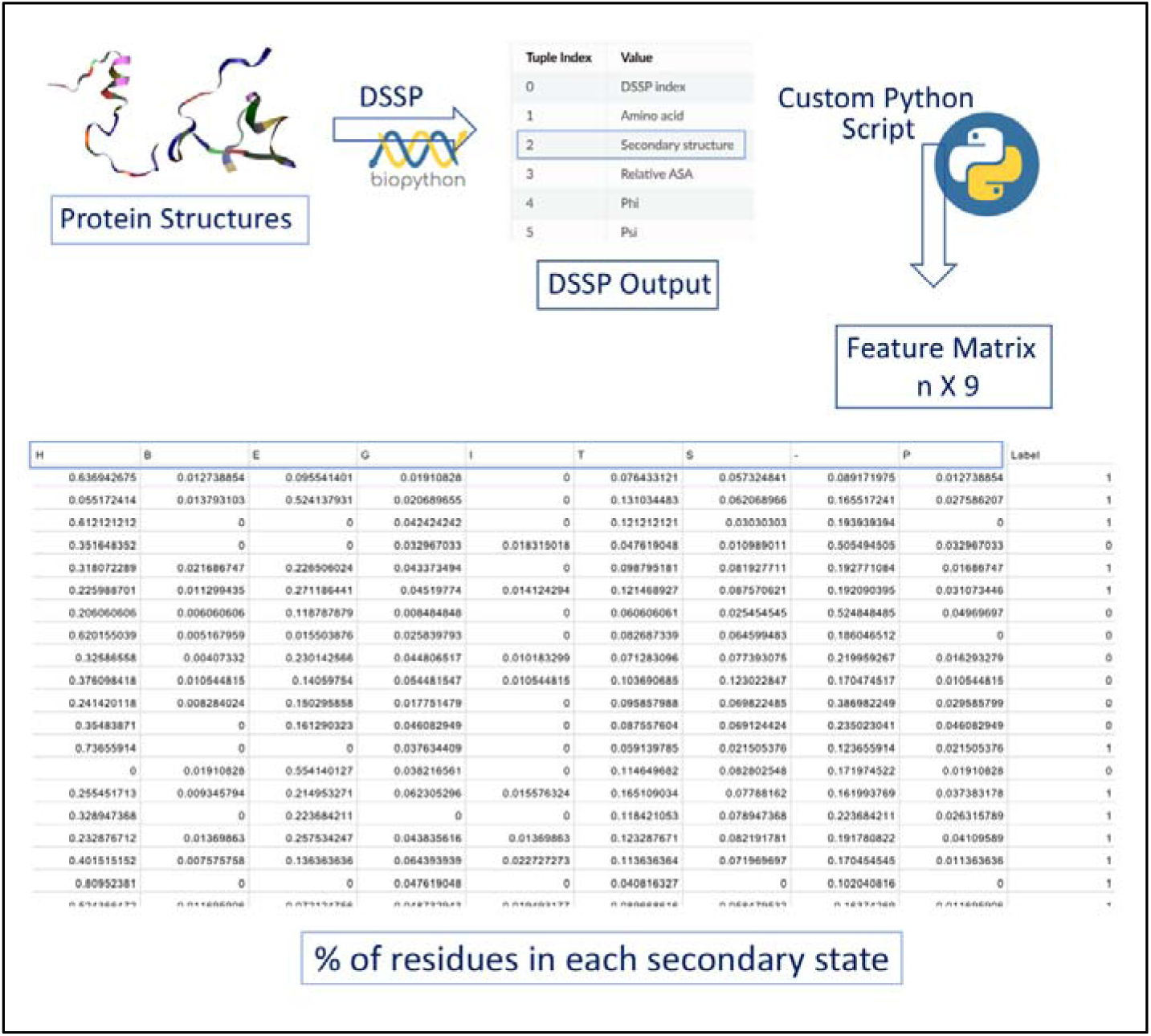
The figure describes the secondary state features in details.

While Relative solvent accessibility (RSA) assign individual residue a unique score based on their extent of burial or exposure of that residue in 3D structure (Tien et al., 2013). We used these values and preprocess them for utilising in prediction task. We have set a threshold based on which we assign each residues in a protein some values such as: for buried cutoff or threshold is 0 <= rsa <= 0.1, for partially burial 0.1< rsa <0.3 and for exposure # rsa < 0.3. Using this approach, we finally calculated 60 features per protein. As the length of proteins are varied in the dataset, calculated RSA values are also vary in terms of ranges. So there is a need of normalizing the dataset. Therefore, we normalize these feature values using standard scaling method and utilize them as RSA features for prediction. Finally, a N*60 size vector is prepared for the RSA based features, for example feature matrix please refer, Figure 3.

**Figure 3:**
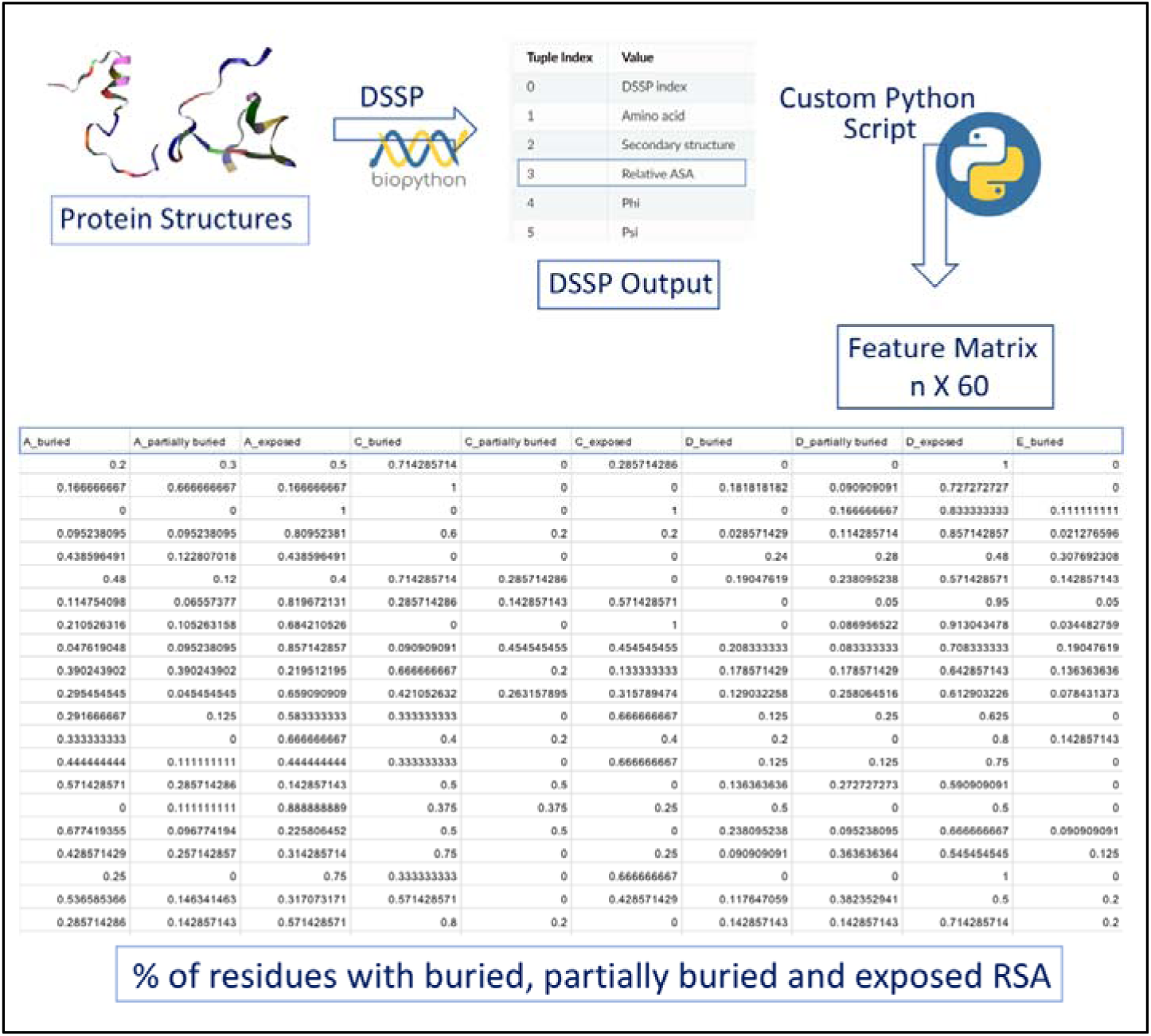
The figure describes the relative solvent accessibility features in details.

### 3.3. Evolutionary profiles based features

#### 3.3.1. PSSM composition based features

It is well recognized that the evolutionary characteristics of the protein offer more important insights into proteins. We have extracted the protein evolutionary information using a position-specific scoring matrix (PSSM) profile generated by Position-Specific Iterated BLAST (PSI-BLAST). For which, we have utilised the “pssm_composition” module from the POSSUM package to construct the PSSM-400, a 20 x 20 dimension vector for a protein sequence composition profile that quantifies the occurrence of 20 amino acids in the sequence. This feature representation effectively summarizes the evolutionary composition of a protein in a compact format.

#### 3.3.2. Raw PSSM profile for Deep Learning based models

In addition to use these compact representations of evolutionary information, use of raw PSSM matrices have also been used in various studies (Khanh Le et al., 2019) that can be used as input features for different DL based models. Since, DL methods do not require handcrafting of features, raw PSSM matrices can be directly used thus retaining the sequential ordering and position-specific evolutionary signals of amino acids. These matrices were treated as input feature maps of size *L×20*, where *L* is the length of the protein sequence and 20 represents the substitution scores for the 20 amino acids at each position. This allowed the deep learning models, particularly convolutional and recurrent layers, to learn spatial patterns and evolutionary motifs across the sequence more effectively. The workflow of DL model is described in the figure 4.

**Figure 4:**
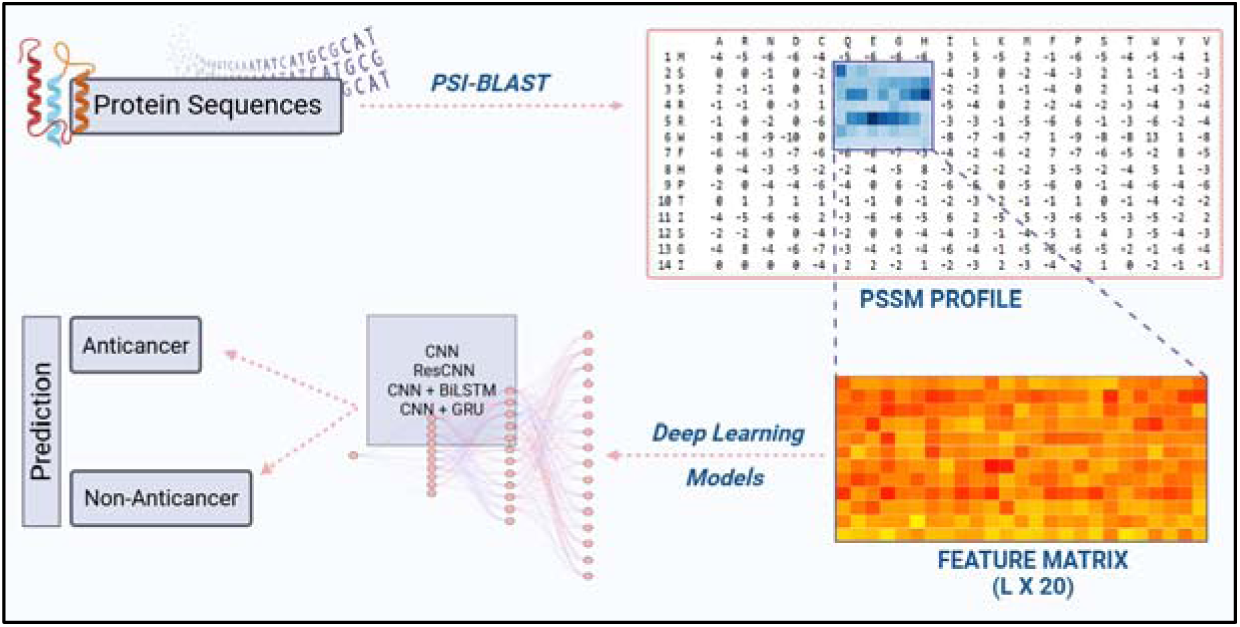
The figure represents the generation of feature matrix using raw PSSM profiles

### 3.4. Sequence embeddings using ESM2

To capture patterns from large-scale multiple sequence alignments (MSAs) implicitly, we have also calculated the embedded features from sequences using 32 layered fine-tuned esm2 model. This model calculated a feature embedding of 1280 dimensional vector.

## 4. Alignment based approach

### 4.1. Basic local alignment search tool (BLAST) for similarity search

Basic Local Alignment Search Tool (BLAST version 2.2.29+) is widely used to annotate protein and nucleotide sequences. Based on a protein’s resemblance to both anticancer and non-anticancer protein sequences, we have used it in this work to identify anti-cancer proteins. The similarity-search based module was developed using the protein–protein BLAST, and the query sequences were compared to the database of anticancer and non-anti-cancer proteins. To do this, anti-cancer proteins were identified using the top hit of BLAST at various E-value cutoffs. Depending on the query sequence’s first database hit, the sequences are designated as either anti-cancer or non-anti-cancer. Various studies have employed this methodology and provided thorough annotations (Boratyn et al., 2012; Kumar et al., 2023; Sharma et al., 2022).

### 4.2. Motif based approach

The Motif-EmeRging and with Classes-Identification (MERCI) tool, a program that can find motifs in any set of sequences, were used to look for the patterns in the anticancer proteins. We have used default parameters of the software to search for recurring patterns found in anticancer protein sequences (Vens et al., 2011).

## 5. Alignment free approach

### 5.1. Machine Learning based classification

Machine learning classifiers are widely used in protein/peptide classification tasks. It provides the best estimators to accurately classify between two classes. In this study, we have developed multiple classification models for distinguishing between anticancer and non-anticancer proteins. The classification models used in this study are Decision Tree (DT), Random Forest (RF), Gradient Boosting (GB), AdaBoost (AB), Extreme Gradient Boost (XGB), Extra Tree (ET), Logistic Regression (LR), K-nearest neighbour (KNN), Support Vector Classifier (SVC), and Multi-layer Perceptron classifier (MLP). We have also optimized these classifiers using various hyperparameters and the best performing models were employed in the study.

### 5.2. Feature Selection Techniques

Prior research has demonstrated that not every feature is significant (Kumar et al., 2025; Teng et al., 2010). As a result, choosing the right features from a wider range of features is quite difficult. To choose the important features from the high-dimensional feature set, we employed Support vector classifier (SVC)-L1-based feature selection (Scikit-learn package) in this study. The SVC-L1 based approach is based on the SVC with linear kernel and penalized with L1 regularization. The L1 regularization forces sparsity by driving less important feature weights to zero, effectively selecting only the most discriminative features and also reduce overfitting. We have enumerated important feature from many combinations using this technique.

### 5.3. Deep Learning based Classification

In this study, we have implemented various deep learning models to classify anticancer proteins using raw evolutionary profiles. Multilayer Perceptron (MLP), Convolutional Neural Network (1D-CNN), Deeper CNN, Residual CNN (ResCNN), CNN + BiGRU (Bidirectional Gated Recurrent Unit) and CNN + BiLSTM (Bidirectional Long Short-Term Memory) have been utilized in this study.

### 5.4. Protein Language Model

To classify anticancer proteins and extract evolutionary features from protein sequences, we used the ESM-2 t33 model (esm2_t33_650M_UR50D), a 650-million-parameter protein language model built on the transformer architecture. For anticancer protein prediction, the model was optimized using Hugging Face Transformers’ EsmForSequenceClassification framework. The base ESM-2 t33 model was initialized with the key hyperparameters such as 33 transformer layers with 20 attention heads per layer, Hidden dimension (hidden_size) = 1280, feedforward dimension (intermediate_size) = 5120, hidden_act=“gelu”, position_embedding_type=“rotary”, layer_norm_eps=0.00001, and initializer_range=0.02.

## 6. Cross-validation and performance evaluation

To overcome the models overfitting/underfitting, we have performed five-fold cross validation over training data. As we have only 205 clusters of 370 anticancer proteins, we manually split our data into five folds so that each fold can have different set of sequences. We have placed each cluster one by one to five folds. The first four clusters were assigned to the training set, while the fifth cluster was used for validation. This process was repeated iteratively until 306 out of the 370 positive sequences were included in the training dataset, with the remaining 64 used for validation. One fold is kept aside for validation purpose while the rest four sets were then again manually splits into five folds to apply cross-validation on training data. This method ensured that the model was evaluated on sequences that were not closely related to those in the training set, thereby minimizing overfitting and enhancing the generalizability of the classifier (**Please refer** figure 2). For negative data, as the sequences retained after CD-HIT were non-redundant by design, they were randomly split into training and validation as same as the number of anticancer proteins present in training and validation sets. In this process, we have randomly select 306 random sequences in training set and 64 sequences to outer validation set. A balanced distribution of positive and negative samples was maintained across both the training and validation subsets, as well as within each fold during cross-validation.

The conventional evaluation measures, such as threshold dependent and independent parameters, were used to evaluate the performance of all machine learning models. Sensitivity, specificity, accuracy, and the Matthews correlation coefficient (MCC) are threshold-dependent characteristics, while the area under the receiver operating characteristic curve (AUC) is threshold-independent.

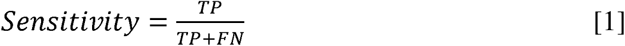

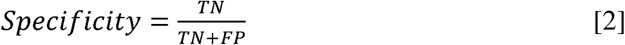

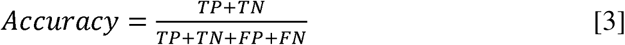

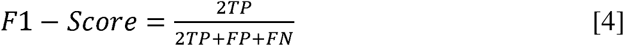

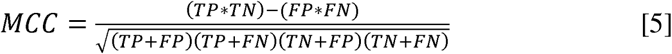

where FP, FN, TP and TN are false positive, false negative, true positive and true negative, respectively.

## 7. Anticancer peptide mapping

This approach is rooted the hypothesis that certain localized regions within a protein sequence might inherently possess functional properties related to anticancer activity. Based on this hypothesis, we retrieved 10mer peptides at different thresholds using protein scan module of AntiCP2 method (Agrawal et al., 2021). This module is designed to develop overlapping protein patterns by choosing the appropriate size. It also predict anticancer properties of these overlapping patterns at different threshold values. Here, we generate 10mer anticancer peptide patterns using each protein sequence and predict them as anticancer at different thresholds [0.15, 0.30, 0.45, 0.60, 0.75, and 0.90]. Using these threshold values, we have got few peptides classified as anticancer. We used number of x obtained peptides at each threshold as feature. Suppose, we have generated 100 peptides of 10mer length from a protein sequence out of which 5 peptides are found to be anticancer at 0.90 threshold, 20 peptides at 0.75, 45 peptides at 0.60, 58 peptides at 0.45, 60 at 0.30 and 70 at 0.15 threshold. Now we use these peptides as feature and numbers as the absolute count of these features. We have also used complete protein length with each peptide as a feature. Also we used a normalized length count calculated by dividing each feature’s absolute count with length of peptide (length of protein is 500 and 5 peptides are extracted from this protein as anticancer, so we divided the 5 by 500 and the obtained ratio is used as normalized count). We have attached an example matrix for detailed understanding of these features in Figure 5.

**Figure 5:**
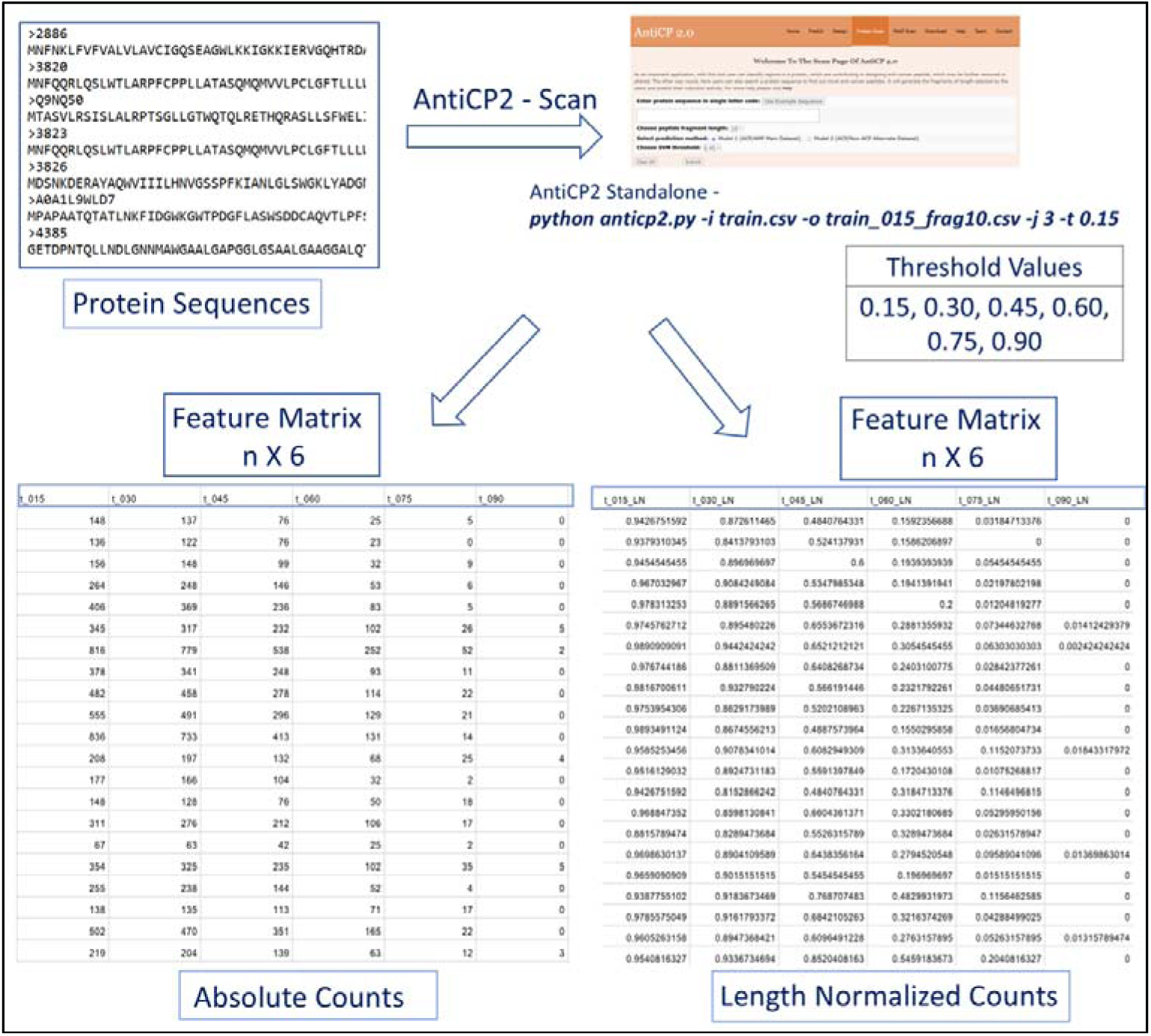
The figure describes the feature matrix prepared using AntiCP2 method’s Protein Scan Module.

**Figure 6:**
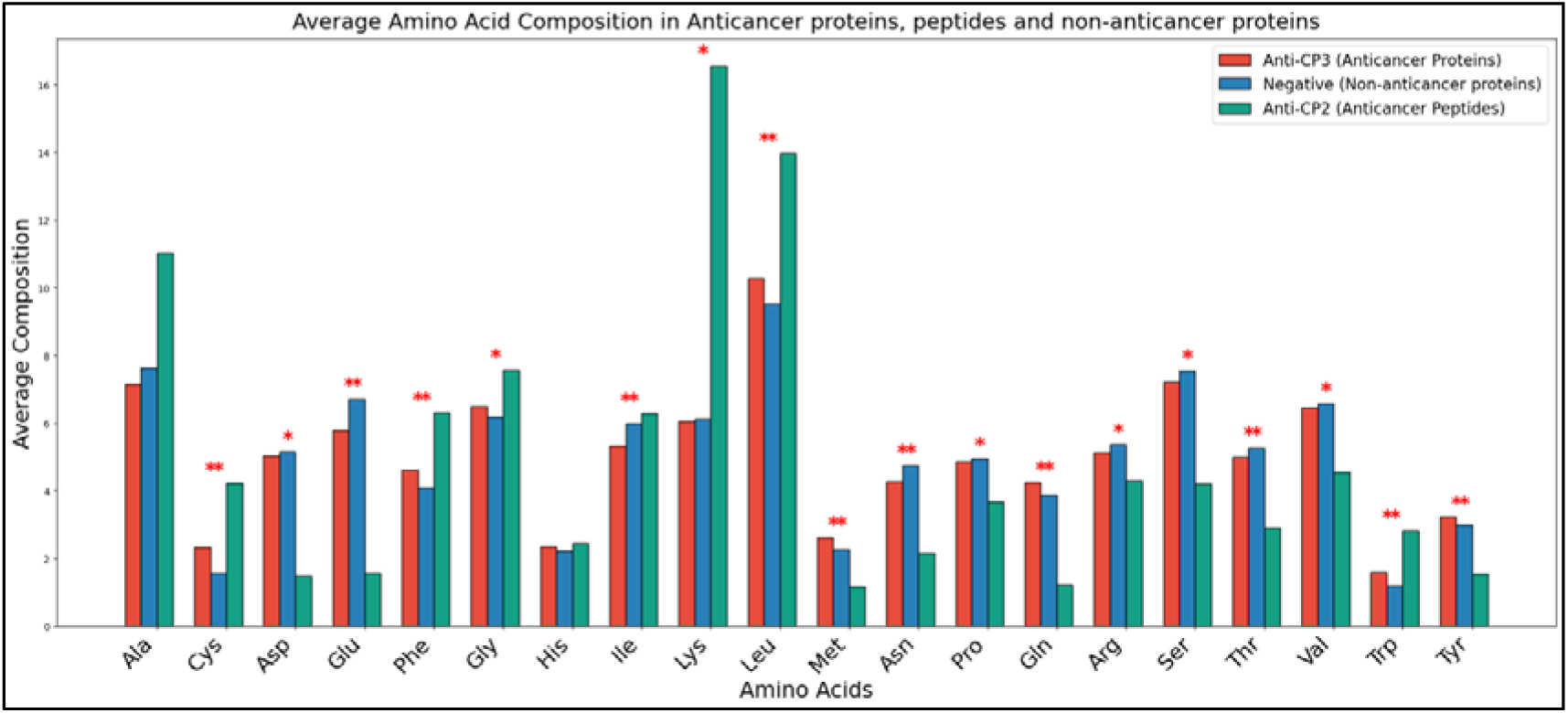
The figure shows the average compositional analysis of amino acids in anticancer proteins, peptides and non-anticancer proteins.

## 8. Hybrid approach

We finally developed a hybrid model which comprises a BLAST based approach with predicted labels of best performing ML classifiers. Merging of these two approaches help to achieve the highest performance and develop a more robust model. To achieve this, firstly BLAST was used to classify the provided protein sequence with an E-value of 10^-20^. We assigned “+0.5” score for positive predictions (anticancer proteins), −0.5 score for negative predictions (non-anticancer proteins and 0 for no hits. When using a hybrid strategy, the total score was calculated by combining the results of two different methodologies (BLAST and ML scores).

## 9. Statistical Analysis

In this study, we have implemented an independent t-test for comparing the average amino acid compositions of anticancer proteins, non-anticancer proteins and anticancer peptides used in AntiCP-2. We have identified significant differences between amino acid residues of all the three datasets based on significant p-value (p<0.05).

## Results

This section provides the detailed analysis and ML performance of different features. Here, we have performed multiple analysis such as composition based analysis, sequence similarity based analysis, conserved regions or Motif based analysis along with ML based analysis using different set of features individually and in combination.

## 1. Compositional Analysis

To identify the type of residues abundant in anticancer proteins, we performed average amino acid compositional analysis between anticancer proteins, peptides and non-anticancer proteins. Independent t-test shows that cysteine, phenylalanine, glycine, leucine and tryptophan residues are significant in anticancer proteins as well as peptides while methionine, glutamine and tyrosine are significantly abundant in only anticancer proteins and lysine is significantly abundant in anticancer peptides only. While aspartic acid, glutamate, asparagine, proline, arginine, serine, threonine, and valine are significantly abundant in non-anticancer proteins. On the basis of compositional analysis study, we can say that the aromatic and non-polar aliphatic amino acids are mostly found in anticancer proteins.

## 2. Alignment-based analysis

### 2.1. BLAST based analysis

With the BLAST-based approach, we have used training data to create a BLAST-formatted database. Next, we compare the training database to query sequences (sequences in the validation set) in order to locate hits at various e-values ranges between 1e^-30^ and 1e^+10^. Depending on whether the top hit was positive or negative, the query sequence assigned some score (for positive score is +0.5, for negative score is −0.5) to classify them as anticancer or non-anticancer. BLAST’s detailed hits on the validation data are displayed in the **Table 1** below.

**Table 1:**
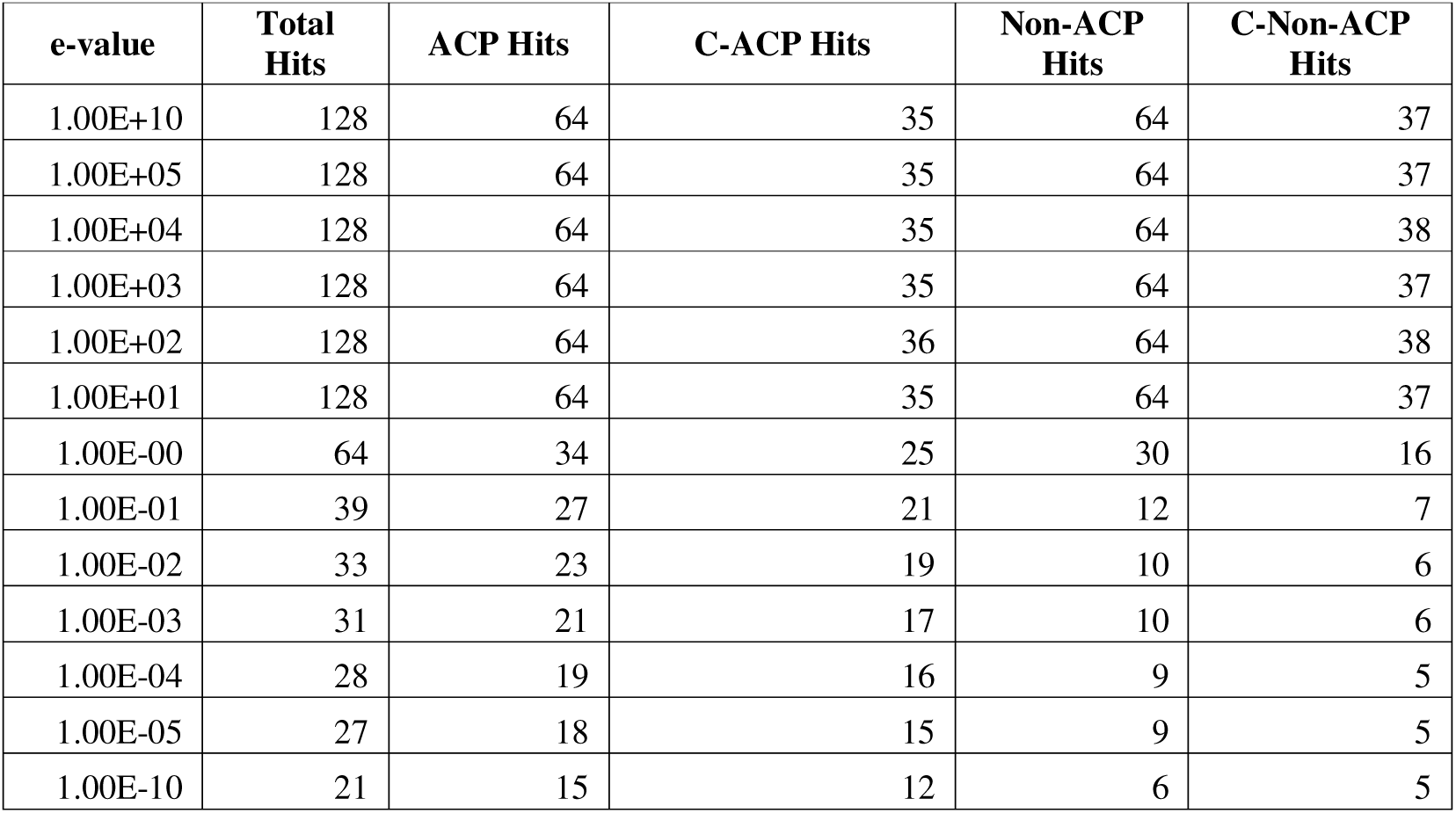
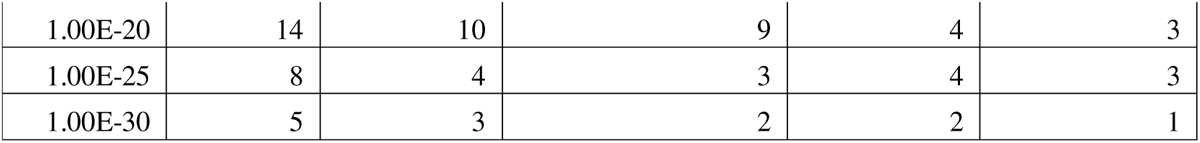
The table describes the BLAST coverage over validation data.

### 2.1. Motif based approach

In motif based approach, we have first identified exclusive anticancer (positive) and non-anticancer (negative) motifs in training data using the MERCI program. Then, we locate these motifs on our validation data. If we found a positive motif in any sequence of validation data, we assign a score to that sequence as +0.5 and if we found a negative motif in any sequence then we assign a score as −0.5 to that sequence. Later, we combine both motif score as well as best ML model’s prediction score together to calculate the other parameter metrices such as accuracy, mcc, area under curve, sensitivity and specificity to evaluate performance. The motif based approach in this data did not perform well (for detailed results, please refer to Supplementary **Table S1**).

## 3. Alignment free analysis

### 3.1. Composition-based features

Several machine learning based classifiers have been applied on composition based features to discriminate between anticancer proteins and non-anticancer proteins. Here, we have calculated the performance over three composition based features namely, amino-acid composition (AAC) based feature, dipeptide composition based feature and physio-chemical property (PCP) composition based feature. The Extra tree (ET) classifier outperform over other classifiers by achieving highest AUROC of 0.69 with MCC of 0.35, 0.30 and 0.28 on amino-acid composition (AAC) based feature, dipeptide composition (DPC) based feature and physio-chemical property composition (PCP) based feature. The AUROC plot of all classifiers on AAC, DPC and PCP features are given below in Figure 7, for detailed results please refer to Supplementary **Table S2.**

**Figure 7:**
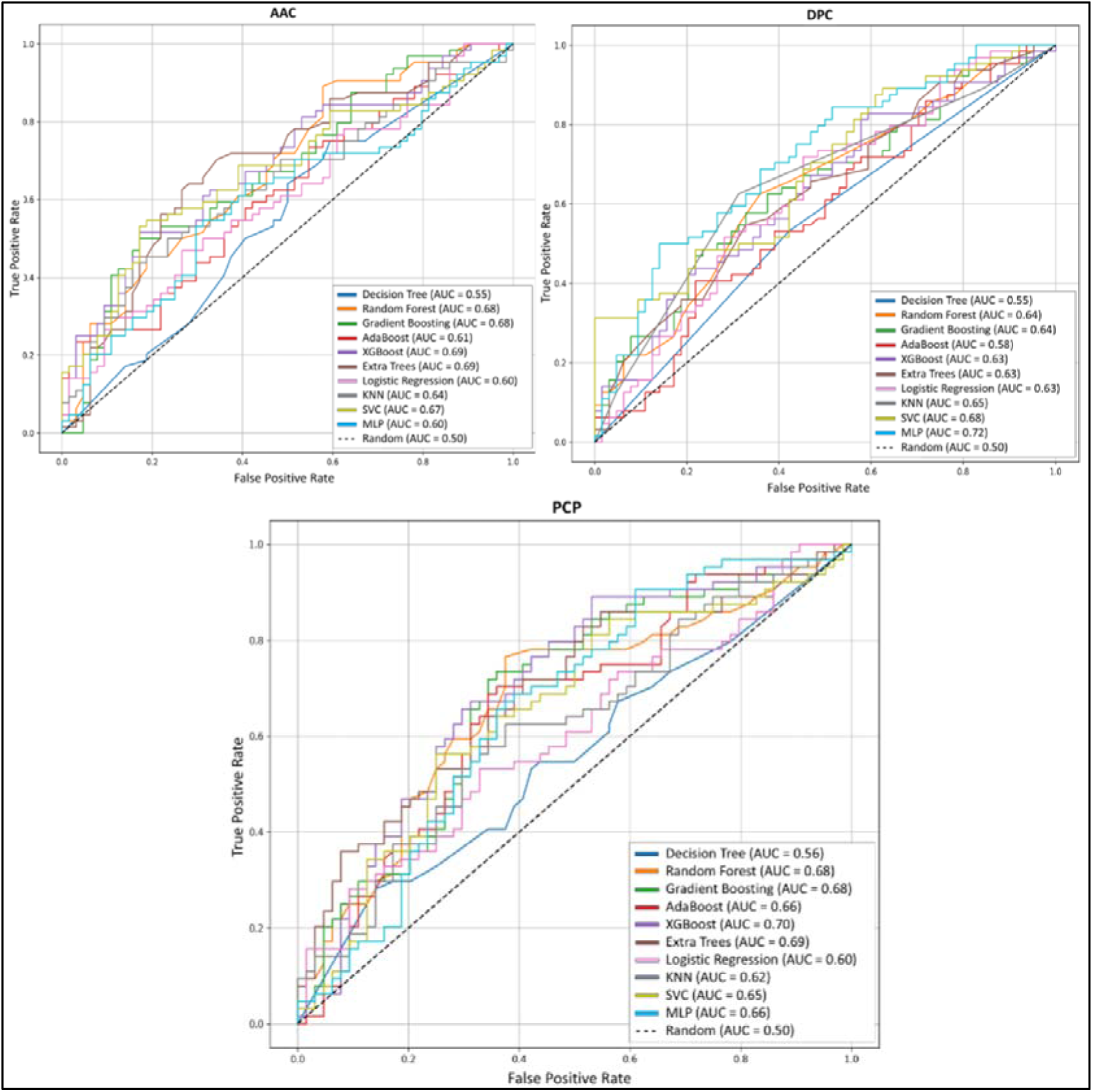
The figure depicts the ML performance of composition based features (AAC, DPC & PCP) over validation dataset.

### 3.2. Secondary structure based features

After calculating and preprocessing secondary structure based features of anticancer proteins. We have applied different machine learning algorithms to classify anticancer proteins based on structure based features. Here, we have observed that MLP classifier outperformed others on RSA features by achieving 0.67 AUC with 0.33 MCC on validation data. While ET outperforms other on Secondary state (DSSP) features by achieving 0.61 AUC on validation data. The AUROC plot of all classifiers on RSA and Secondary state features are given below in Figure 8. The detailed results of Secondary structure based features are given in **Supplementary Table S3**.

**Figure 8:**
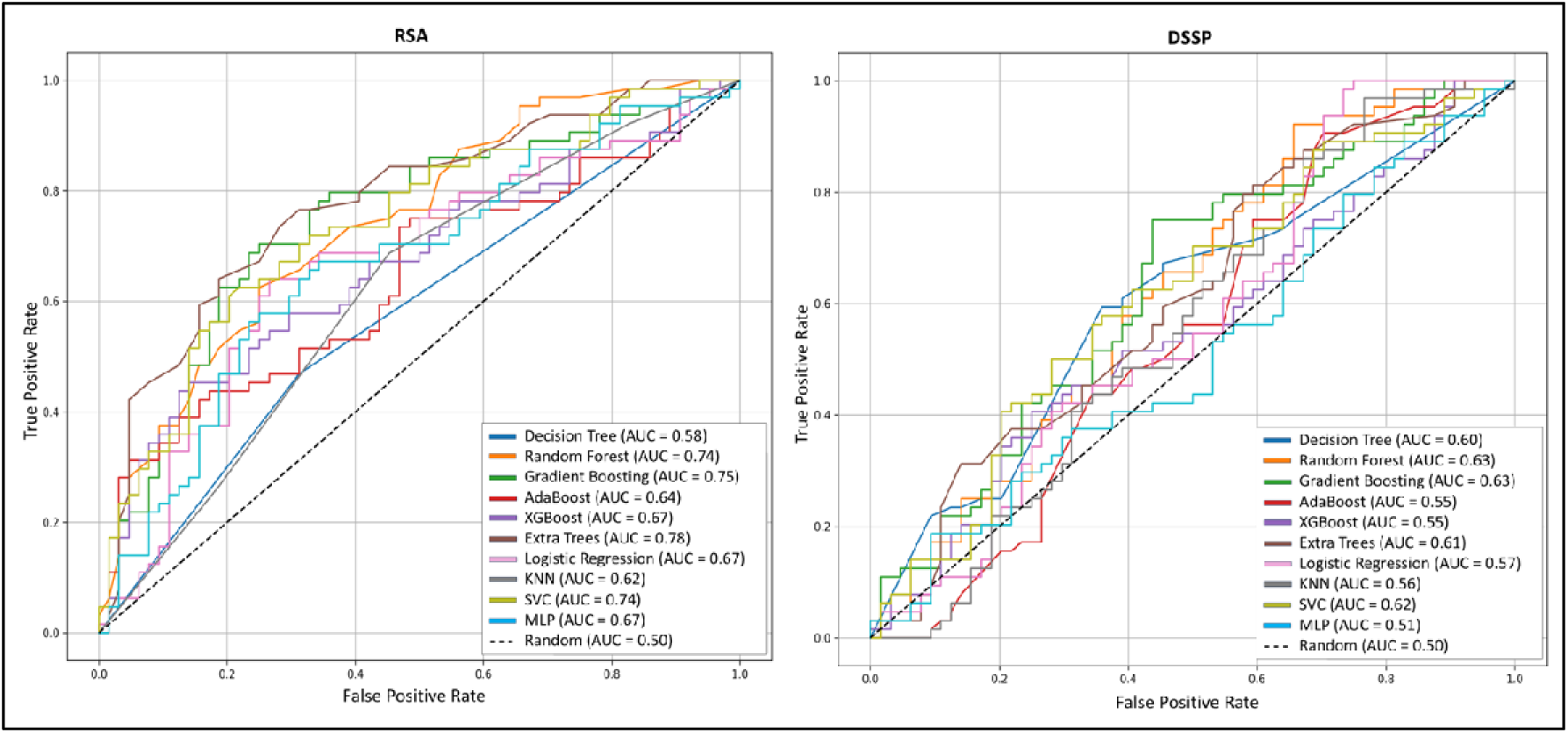
The figure shows the ML performance on secondary structure based features over validation data.

### 3.3. Evolutionary profile based features

#### 3.3.1. PSSM composition based features

We have also developed a model based on PSSM profiles, a 20 x 20 dimension vector for a protein sequence composition profile to classify anticancer proteins. In this case, the MLP classifier outperforms other classifiers and achieves maximum AUROC of 0.79 and MCC of 0.44 on validation dataset. The evolutionary profile based feature outperforms other types of features. The AUROC plot of all classifiers on PSSM-composition based features are given below in Figure 9, for detailed results please refer to Supplementary **Table S4.**

**Figure 9:**
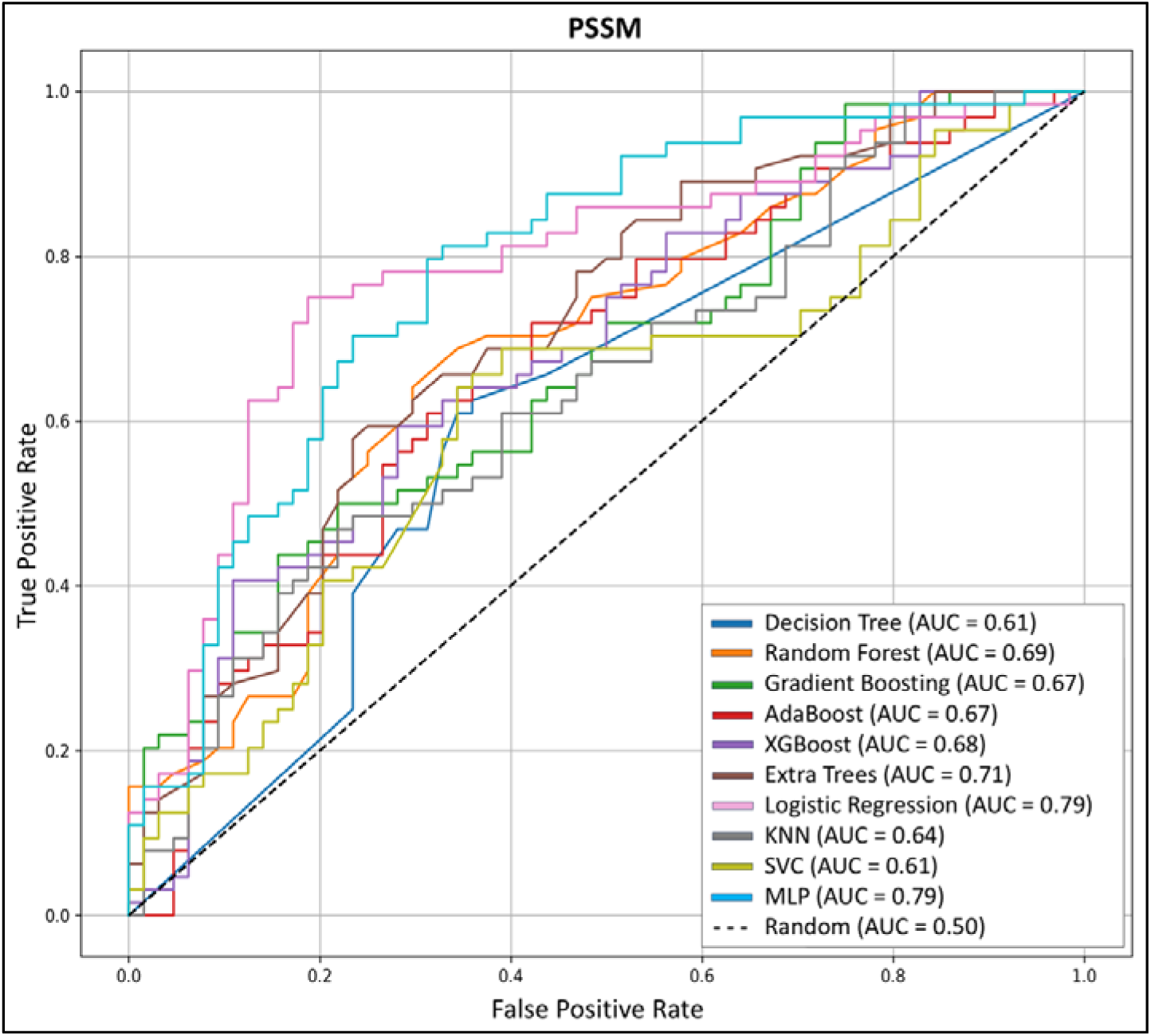
The figure shows the ML performance on evolutionary profile (PSSM) based features on validation dataset.

#### 3.3.2. Raw PSSM profile for Deep Learning based models

We have explored a series of Deep Learning architectures trained directly on the raw PSSM profiles. While the DL models show potential in capturing rich local and sequential dependencies, the traditional machine learning approach using PSSM-composition features still outperforms the DL models in terms of overall accuracy and robustness. One of the main limitations that we have in developing these models is limited data availability. Deep learning models typically require large amounts of data to generalize well. The detailed results of implemented DL models can be seen in **Table 2**.

**Table 2:** The table shows the performance of DL models implemented on raw PSSM profiles.

| Model | Sens | Spec | Acc | AUC | MCC |
| --- | --- | --- | --- | --- | --- |
| MLP (2 Dense Layers, Dropout) | 0.46 | 0.7 | 0.58 | 0.61 | 0.17 |
| **CNN(Conv1D + MaxPool)** | 0.59 | 0.75 | 0.67 | 0.7 | 0.34 |
| **Deeper CNN (3 Conv1D + MaxPool)** | 0.62 | 0.61 | 0.61 | 0.67 | 0.23 |
| **ResCNN(CNN with Residual Blocks)** | 0.59 | 0.75 | 0.67 | 0.74 | 0.34 |
| **CNN+BiGRU(1 Conv1D + BiGRU)** | 0.46 | 0.78 | 0.62 | 0.7 | 0.26 |
| **CNN+BiLSTM((1 Conv1D + BiLSTM)** | 0.54 | 0.75 | 0.65 | 0.72 | 0.30 |

### 3.4. SVC-L1 based Feature selection

We have applied well-known feature selection technique: SVC-L1 on different features as well as on combination of features. The list of number of selected features from set of individual features as well as combined feature pool is given in **Supplementary Table S5**. Here, we have retrieve a set of 256 best features from PSSM composition based features. We have applied different ML-based classifiers on these selected features and found that the MLP classifier achieved the highest AUROC of 0.78 and MCC of 0.42 on validation dataset. The detailed results of multiple classifiers on 256 features is given below, see **Table 3**. The detailed results over different combinations, please refer **Supplementary S6**.

**Table 3:** The table shows the performance of set of PSSM features selected using SVC-L1 method.

| Model | Sensitivity | Specificity | Accuracy | AUC | MCC |
| --- | --- | --- | --- | --- | --- |
| **Decision Tree** | 32.81 | 73.44 | 53.12 | 0.58 | 0.07 |
| **Random Forest** | 57.81 | 67.19 | 62.5 | 0.7 | 0.25 |
| **Gradient Boosting** | 50 | 78.12 | 64.06 | 0.69 | 0.29 |
| **AdaBoost** | 37.5 | 75 | 56.25 | 0.63 | 0.13 |
| **XGBoost** | 48.44 | 81.25 | 64.84 | 0.7 | 0.31 |
| **Extra Trees** | 54.69 | 79.69 | 67.19 | 0.73 | 0.36 |
| **Logistic Regression** | 56.25 | 82.81 | 69.53 | 0.78 | 0.41 |
| **KNN** | 53.12 | 64.06 | 58.59 | 0.64 | 0.17 |
| **SVC** | 100 | 0 | 50 | 0.39 | 0 |
| **MLP** | 57.81 | 82.81 | 70.31 | 0.78 | 0.42 |

### 3.5. Protein Language Model

We have used different layers of esm2 model for classification of anticancer proteins such as t6 layer, t12 layer, t30 layer and t33layer. After finetuning this model, the highest AUC (0.90) was achieved by ESM2-t33 layer with MCC of 0.63. The detailed results of ESM2 model using different layers are given in Table 4.

**Table 4:**
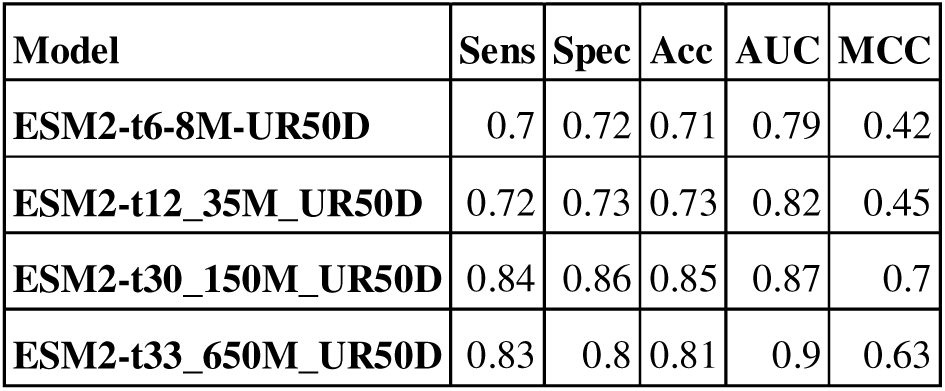
The table shows the performance of fine-tuned ESM2 model.

| Model | Sens | Spec | Acc | AUC | MCC |
| --- | --- | --- | --- | --- | --- |
| ESM2-t6-8M-UR50D | 0.7 | 0.72 | 0.71 | 0.79 | 0.42 |
| ESM2-t12_35M_UR50D | 0.72 | 0.73 | 0.73 | 0.82 | 0.45 |
| ESM2-t30_150M_UR50D | 0.84 | 0.86 | 0.85 | 0.87 | 0.7 |
| ESM2-t33_650M_UR50D | 0.83 | 0.8 | 0.81 | 0.9 | 0.63 |

Furthermore, since embeddings generated by the fine-tuned model can serve as valuable features, we have extracted embeddings using the above finetune model and applied various ML algorithms. The finetune model also achieved highest AUC 0.90 which is same as of finetune model, refer result **Table 5**. We have incorporated the best ESM2 finetuned model in webserver and standalone versions.

**Table 5:**
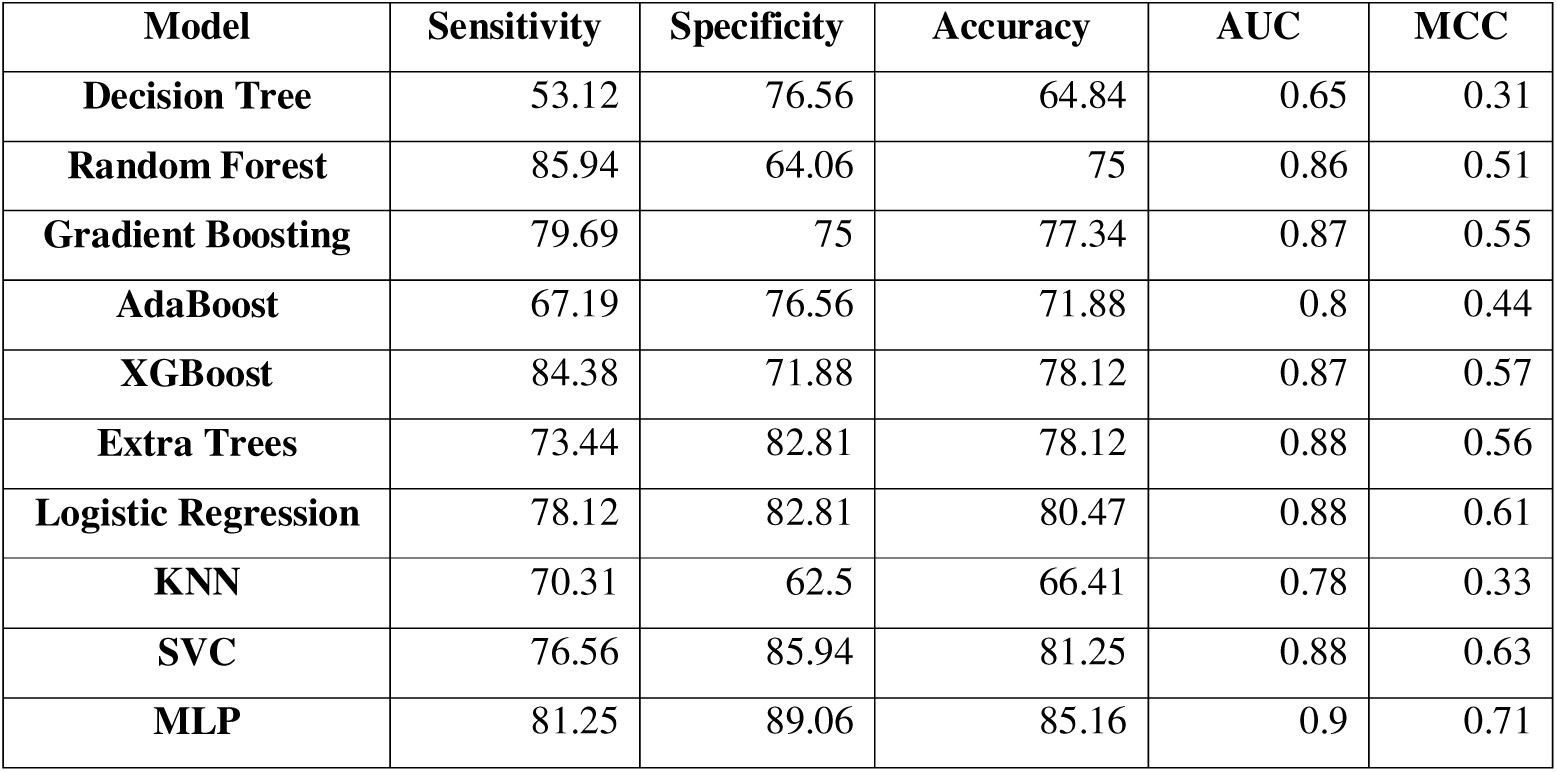
The table shows the ML performance of embeddings extracted using ESM2-t33 model.

| Model | Sensitivity | Specificity | Accuracy | AUC | MCC |
| --- | --- | --- | --- | --- | --- |
| Decision Tree | 53.12 | 76.56 | 64.84 | 0.65 | 0.31 |
| Random Forest | 85.94 | 64.06 | 75 | 0.86 | 0.51 |
| Gradient Boosting | 79.69 | 75 | 77.34 | 0.87 | 0.55 |
| AdaBoost | 67.19 | 76.56 | 71.88 | 0.8 | 0.44 |
| XGBoost | 84.38 | 71.88 | 78.12 | 0.87 | 0.57 |
| Extra Trees | 73.44 | 82.81 | 78.12 | 0.88 | 0.56 |
| Logistic Regression | 78.12 | 82.81 | 80.47 | 0.88 | 0.61 |
| KNN | 70.31 | 62.5 | 66.41 | 0.78 | 0.33 |
| SVC | 76.56 | 85.94 | 81.25 | 0.88 | 0.63 |
| MLP | 81.25 | 89.06 | 85.16 | 0.9 | 0.71 |

## 4. Combined feature evaluation

In order to identify for more relevant feature, we have also tried different combinations of features. In these combination, we have observed that PSSM and DSSP combination performs much better than other combinations and achieved AUROC of 0.81 with MCC of 0.49. The list of combinations with their performance are given below in **Table 6**, for detailed results please refer **Supplementary table S7.**

**Table 6:** The table shows the performance over different combined features using ML classifiers.

| **Feature Type** | **Model Name** | **Sens** | **Spec** | **Acc** | **AUC** | **MCC** |
| --- | --- | --- | --- | --- | --- | --- |
| **AAC + DPC** | Extra Trees | 65.62 | 67.19 | 66.41 | 0.71 | 0.33 |
| **AAC + PCP** | Gradient Boosting | 68.75 | 60.94 | 64.84 | 0.7 | 0.3 |
| **AAC + DSSP** | Extra Trees | 76.56 | 68.75 | 72.66 | 0.77 | 0.45 |
| **AAC + RSA** | Gradient Boosting | 64.06 | 75 | 69.53 | 0.71 | 0.39 |
| **AAC + PSSM** | Logistic Regression | 62.5 | 87.5 | 75 | 0.78 | 0.52 |
| **AAC + DSSP + RSA** | Extra Trees | 73.44 | 70.31 | 71.88 | 0.79 | 0.44 |
| **DSSP + RSA** | Random Forest | 56.25 | 82.81 | 69.53 | 0.77 | 0.41 |
| **DPC + PCP** | Extra Trees | 71.88 | 71.88 | 71.88 | 0.75 | 0.44 |
| **DPC + RSA** | Random Forest | 68.75 | 81.25 | 75 | 0.79 | 0.5 |
| **DPC + DSSP** | Random Forest | 64.06 | 65.62 | 64.84 | 0.73 | 0.3 |
| **DPC + DSSP + RSA** | Extra Trees | 60.94 | 78.12 | 69.53 | 0.76 | 0.4 |
| **DPC + PSSM** | Random Forest | 62.5 | 75 | 68.75 | 0.71 | 0.38 |
| **PCP + PSSM** | Logistic Regression | 62.5 | 84.38 | 73.44 | 0.78 | 0.48 |
| **PCP + DSSP** | Gradient Boosting | 68.75 | 71.88 | 70.31 | 0.74 | 0.41 |
| **PCP + RSA** | AdaBoost | 60.94 | 78.12 | 69.53 | 0.72 | 0.4 |
| **PCP +**<br>**DSSP +**<br>**RSA** | Extra Trees | 82.81 | 62.5 | 72.66 | 0.79 | 0.46 |
| **PSSM +**<br>**DSSP** | Logistic<br>Regression | 67.19 | 81.25 | 74.22 | 0.81 | 0.49 |
| **PSSM +**<br>**RSA** | Logistic<br>Regression | 64.06 | 82.81 | 73.44 | 0.78 | 0.48 |
| **PSSM +**<br>**DSSP +**<br>**RSA** | Logistic<br>Regression | 64.06 | 82.81 | 73.44 | 0.8 | 0.48 |

## 5. Anticancer peptide mapping

We hypothesised that the short peptide segments designed from whole proteins can capture a more granular representation of the protein’s potential anticancer characteristics. In this study, we tried different peptide patterns at multiple threshold values as features. Here, we have found that a combined set of 7 features including direct count (6 features), peptide length (1 feature) and 13 features from the combined set of direct count (6 features), peptide length (1 feature) and normalized count (6 features) obtained 0.58 AUC performance on the validation set by applying extreme gradient boosting (XGB) and gradient boosting (GB) classifier (**see Table 7**). Based on the performance achieved on these 10-mers, we comes to the conclusion that this hypothesis is not worked here. One potential reason for this could be the limitations of the 10-mer-based approach itself. Anticancer activity might not be dictated solely by the properties of individual 10-mers but instead by complex structural or sequence-level interactions that extend beyond short stretches. Additionally, AntiCP2 was trained on shorter peptides (length <= 40), so the tool may not be able accurately capture the nuanced relationships between local peptide properties and overall protein activity. These factors suggest that while this approach provided an interesting perspective, further exploration may be necessary for improving the predictive performance. For detailed results over other set of features, please refer **Supplementary Table S8**.

**Table 7:** The table shows the ML performance of different short peptide features over independent dataset.

| Feature | Model | Sensitivity | Specificity | Accuracy | AUC | MCC |
| --- | --- | --- | --- | --- | --- | --- |
| AntiCP2 - DC | ET | 54.69 | 54.69 | 54.69 | 0.55 | 0.09 |
| AntiCP2 - LN | DT | 65.62 | 46.88 | 56.25 | 0.58 | 0.13 |
| AntiCP2 – DC +<br>Length | XGBoost | 53.12 | 65.62 | 59.38 | 0.58 | 0.19 |
| AntiCP2 – DC +<br>Length + LN | GB | 51.56 | 56.25 | 53.91 | 0.56 | 0.08 |

## 6. Hybrid approach

To develop a more accurate model, we combined alignment based approach (BLAST) with alignment free approach (ML). Here, we combined the best e-value (10e^-20^) hit BLAST scores with ML scores of best performing LLM model. Finally, we achieve highest AUC of 0.91 and MCC of 0.63 on validation dataset using the combined scores. The detailed results can be seen below in **Table 8**.

**Table 8:** The table shows the performance of best hybrid model developed using BLAST and top selected features.

| e-value | Sens | Spec | Accuracy | AUC | Kappa | MCC |
| --- | --- | --- | --- | --- | --- | --- |
| 1.00E+01 | 54.69 | 57.81 | 56.25 | 0.76 | 0.12 | 0.13 |
| 1.00E+02 | 56.25 | 59.38 | 57.81 | 0.77 | 0.16 | 0.16 |
| 1.00E+03 | 54.69 | 57.81 | 56.25 | 0.76 | 0.12 | 0.13 |
| 1.00E+04 | 54.69 | 59.38 | 57.03 | 0.77 | 0.14 | 0.14 |
| 1.00E+05 | 54.69 | 57.81 | 56.25 | 0.76 | 0.12 | 0.13 |
| 1.00E+10 | 54.69 | 57.81 | 56.25 | 0.76 | 0.12 | 0.13 |
| 0 | 78.12 | 65.62 | 71.88 | 0.85 | 0.44 | 0.44 |
| 1.00E-01 | 79.69 | 75 | 77.34 | 0.89 | 0.55 | 0.55 |
| 1.00E-02 | 82.81 | 75 | 78.91 | 0.9 | 0.58 | 0.58 |
| 1.00E-03 | 82.81 | 75 | 78.91 | 0.9 | 0.58 | 0.58 |
| 1.00E-04 | 82.81 | 75 | 78.91 | 0.9 | 0.58 | 0.58 |
| 1.00E-05 | 82.81 | 75 | 78.91 | 0.9 | 0.58 | 0.58 |
| 1.00E-10 | 82.81 | 79.69 | 81.25 | 0.9 | 0.62 | 0.63 |
| 1.00E-20 | 84.38 | 78.12 | 81.25 | 0.91 | 0.62 | 0.63 |

## 7. Design and implementation of web server

We have made a systematic attempt to develop a web-based application for prediction of anticancer proteins. We have incorporated our best performing models: Model 1 developed using esm2-t33 model and Model 2 developed using hybrid approach with combination of BLAST and LLM model. We have developed two major modules in the webserver. The Predict module includes two models, first model is completely based on ML-based classification and another model is based on hybrid approach which includes BLAST and LLM based classification. We have also developed another module named as BLAST module which is based on sequence similarity search. The detailed architecture of the web server is shown below in Figure 10. We have also developed standalone, GitHub and pip package for this method. The anticp3 web server is available at https://webs.iiitd.edu.in/raghava/anticp3, standalone at https://webs.iiitd.edu.in/raghava/anticp3/down.html, GitHub at https://github.com/raghavagps/anticp3, pip package at https://pypi.org/project/anticp3/ and hugging face https://huggingface.co/raghavagps-group/anticp3.

**Figure 10:**
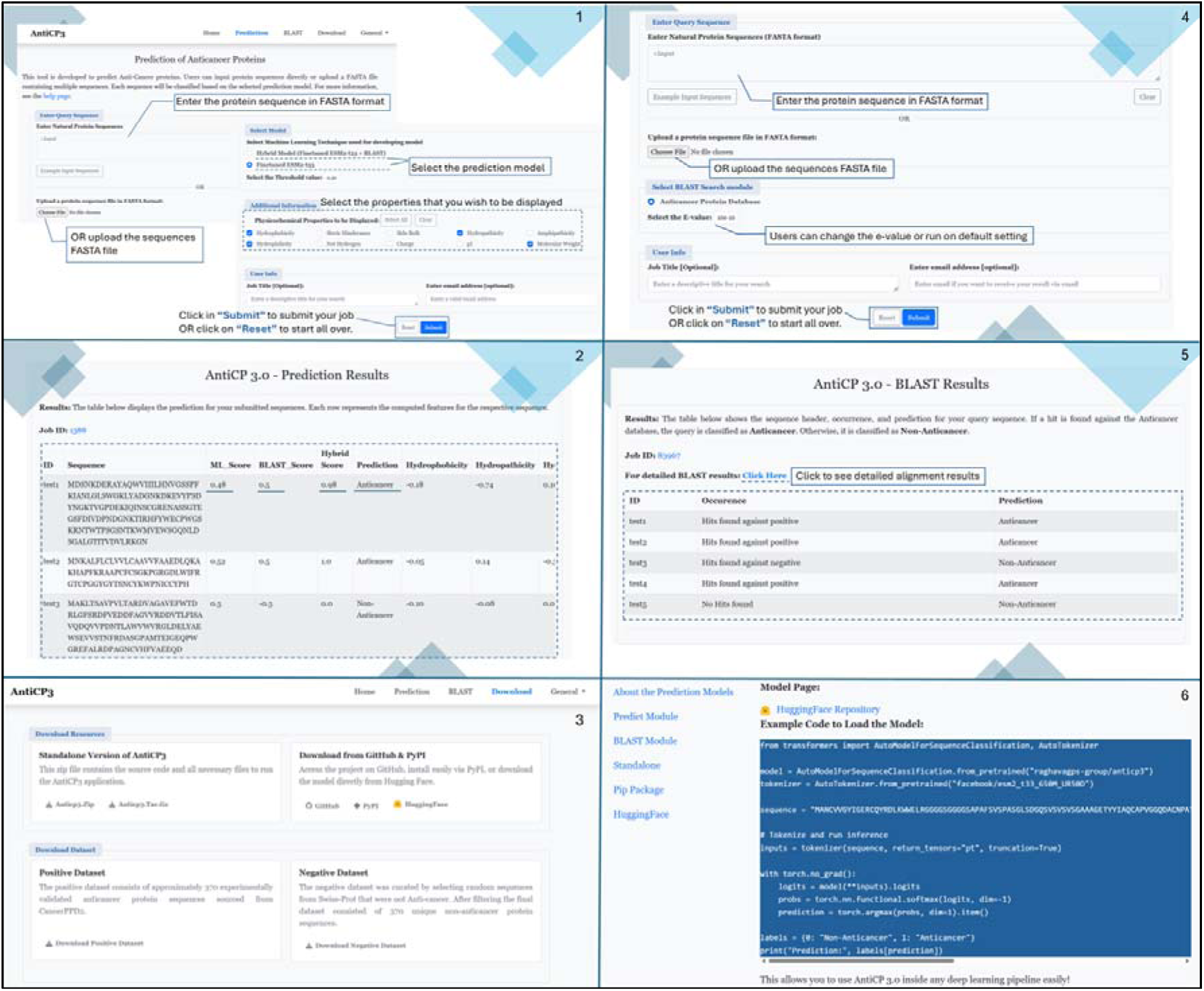
The figure represents the architecture of AntiCP3 webserver.

## 8. Comparison with other method

The AntiCP3 webserver is developed using anticancer protein sequences with length more than 50. Previously, multiple attempts were made to develop peptide based methods for anticancer peptide classification. The first well-known method Anti-CP published in 2013, which was developed using 225 anticancer peptides sequences with peptide length of less than 50 (Tyagi et al., 2013). After this method, several other methods were developed such as Hajisharifi et al; mACPred, TargetACP, ACPred, AntiCP2.0, ACP-ML, MLACP-2.0 and many more (Agrawal et al., 2021; Boopathi et al., 2019; Hajisharifi et al., 2014; Schaduangrat et al., 2019; Thi Phan et al., 2022). But all methods were designed for the classification of anticancer peptide. We have found two methods whose webserver accept protein sequences although they have mentioned in paper that their model was trained on peptide sequences. The first method is CancerGram, they have mentioned that CancerGram model was trained on peptide dataset but their web server accepts protein sequences as well (Burdukiewicz et al., 2020). We have classified our validation data on CancerGram web server, this webserver predicts all protein sequences as negative and provides AUC 0.52. The another method is mACPred 2.0 (Sangaraju et al., 2024), this method performed better than CancerGram over our validation dataset and achieves 0.66 AUC. Till Now, there is no method developed for anticancer protein classification. This is the first attempt where complete protein sequences were used for model training to distinguish between anticancer proteins from non-anticancer proteins.

## Discussion & Conclusion

Anticancer protein plays an important role in the therapeutic purposes of cancer. Multiple drugs and therapies are available for the cancer treatment such as monoclonal antibodies (mABs), small-molecule drugs, and cancer growth blockers (Mukherjee et al., 2023). It is not necessary that the peptide/protein can have only one property. There are many proteins/peptides that are antimicrobial in nature but also exhibit anticancer properties. Similarly, multiple anticancer antibiotics prepared by extracting proteins from bacteria’s having anticancer properties such as Bleomycin retrieved from *Streptomyces verticillus* targets head and neck squamous cell carcinomas, Hodgkin’s disease, non-Hodgkin’s lymphoma, testicular carcinomas, ovarian cancer and malignant pleural effusion (Karpinski & Adamczak, 2018). As identification and screening of potential anticancer proteins in the wet lab is time-consuming, costly and labour-intensive process, there is a need for in silico tools that can predict anticancer proteins with high reliability. Previously, various studies have been designed for the classification and prediction of anticancer peptides. For a detailed list of prediction methods for peptides, please refer to Supplementary **Table S9**. In this study, we are majorly focusing on the prediction and classification of anticancer proteins as they have their own importance over peptides.

Here, we analysed different types of features of anticancer proteins such as composition based feature, evolutionary profile based, secondary structure based and sequence derived features. Each feature has its own importance as composition based features provide insights about specific arrangement of amino acids within a sequence while the evolutionary profile of protein provides the amino acid positions having highest predictive power for a certain biological feature (Harding-Larsen et al., 2024). We have observed that as all types of features work well for classifying between anticancer proteins from non-anticancer proteins, only anticancer mapping peptide features have the least performance in comparison to others. This can happen due to two major reasons, 1. Complex structural or sequence-level interactions that extend across short lengths may determine anticancer activity rather than just the characteristics of individual 10-mers. 2. AntiCP2 may not be able to precisely capture the complex interactions between local peptide characteristics and overall protein activity because it was trained on shorter peptides (length <= 40). We have also tried different feature combinations and also performed feature selection techniques over these combinations. The ESM2 based fine-tuned model outperforms other classifiers and achieved a highest AUC of 0.90 with MCC as 0.71 on validation data. Using these features we have developed a hybrid model in combination with BLAST to increase the models performance to 0.91 as AUC and 0.63 as MCC. We have also provided the standalone, GitHub and pypi package along with webserver by incorporating our best performing models. We anticipate that this approach will accelerate the search for and development of innovative, potent anticancer protein-based treatments.

## Acknowledgements

Authors are thankful to, All India Council for Technical Education (AICTE), Department of Science and Technology (DST-INSPIRE), Indraprastha Institute of Information Technology (IIITD), for fellowships and the financial support and Department of Computational Biology, IIITD, New Delhi, for infrastructure and facilities. We thank the DBT for providing an infrastructure grant to the institute.

## Availability

The webserver AntiCP3 is freely accessible at https://webs.iiitd.edu.in/raghava/anticp3/”. Dataset and standalone version can be downloaded from the downloads tab given in the website at https://webs.iiitd.edu.in/raghava/anticp3/down.html. The GitHub, pypi and hugging face packages are available at https://github.com/raghavagps/anticp3, https://pypi.org/project/anticp3/ and https://huggingface.co/raghavagps-group/anticp3.

## Authors Contribution

A.G., M.C., R.T., and G.P.S.R. collected and processed the datasets. A.G., M.C., R.T., and G.P.S.R. implemented the algorithms and developed the prediction models. A.G., M.C., R.T., and G.P.S.R. analysed the results. A.G., and M.C., created the front-end user interface and back end of the webserver. R.T., and G.P.S.R. penned the manuscript. R.T., M.C., A.G. and G.P.S.R. reviewed the manuscript. G.P.S.R. conceived and coordinated the project. All authors read and approved the final manuscript.

